# Dopamine facilitates fear memory formation in the human amygdala

**DOI:** 10.1101/2021.03.31.437765

**Authors:** Andreas Frick, Johannes Björkstrand, Mark Lubberink, Allison Eriksson, Mats Fredrikson, Fredrik Åhs

## Abstract

Learning which environmental cues that predict danger is crucial for survival and accomplished through Pavlovian fear conditioning. In humans and rodents alike, fear conditioning is amygdala-dependent and rests on similar neurocircuitry. Rodent studies have implicated a causative role for dopamine in the amygdala during fear memory formation, but the role of dopamine in aversive learning in humans is unclear. Here, we show dopamine release in the amygdala and striatum during fear learning in humans. Using simultaneous positron emission tomography and functional magnetic resonance imaging, we demonstrate that the amount of dopamine release is linked to strength of conditioned fear responses and linearly coupled to learning-induced memory trace activity in the amygdala. Thus, like in rodents, formation of amygdala-dependent fear memories in humans seems to be facilitated by endogenous dopamine release, supporting an evolutionary conserved neurochemical mechanism for aversive memory formation.

---

Fear conditioning is an evolutionary shaped mechanism for aversive memory formation important for survival. In fear conditioning, a previously neutral cue turns into a conditioned stimulus (CS) through pairings with an aversive stimulus (*1*), forming a memory trace in the amygdala (*2–4*), a key brain region supporting associative and emotional learning (*5, 6*). The amygdala is heavily innervated by dopamine (*7*), and in rodents, optogenetic stimulation of dopaminergic neurons (*8*), as well as systemic and amygdala-targeted administration of dopaminergic agonists, increase dopamine signaling and facilitate aversive learning (*9*). In contrast, dopamine antagonists lead to attenuated memory formation (*10*). Similarly, neurochemical lesions to the dopamine system in the amygdala severely compromise fear learning (*11, 12*), and fear conditioning does not occur in genetically dopamine-deficient mice, but is restored with administration of the dopamine precursor _L_-DOPA (*13*). Dopamine release seems both necessary (*13*) and sufficient (*14*) for fear conditioning in rodents, suggesting causation and implicating that human fear learning is dopamine-dependent. However, little is known of dopaminergic modulation of aversive learning in humans, but working memory (*15, 16*) and sequential learning (*17*) are facilitated by endogenous dopamine release in the striatum. In Parkinson patients with reduced dopamine function, amygdala-mediated fear processing is compromised but restored after dopamimetic treatment (*18*), and polymorphisms in genes encoding dopamine receptor 4 are associated with human fear conditioning (*19*). However, no brain imaging study has directly evaluated if dopamine is released during amygdala-mediated associative learning or if the amount of dopamine released predicts learning strength.

To assess if fear memory formation in humans is dopamine-related, we simultaneously measured brain dopamine release and neural activity in a combined positron emission tomography/magnetic resonance imaging (PET/MRI) scanner during fear conditioning, with fear learning probed by skin conductance responses (SCR) to a shock-predicting cue (CS+) and a control cue (CS-) never paired with shock (Fig. 1). We used single scan bolus/infusion of [^11^C]raclopride (*20, 21*) to measure conditioning-related change in binding potential (i.e., dopamine release) (*15*), combined with functional magnetic resonance imaging (fMRI) to measure neural activity. We predicted that increased dopamine levels in the amygdala during fear conditioning would facilitate learning (*8, 13, 14*), with greater dopamine release related to superior cue discrimination and enhanced amygdala-located fear engram formation. We also conducted exploratory analyses in the striatum, another brain region involved in fear conditioning rich in dopamine D2/3 receptors (*22*).

**Fig. 1.**
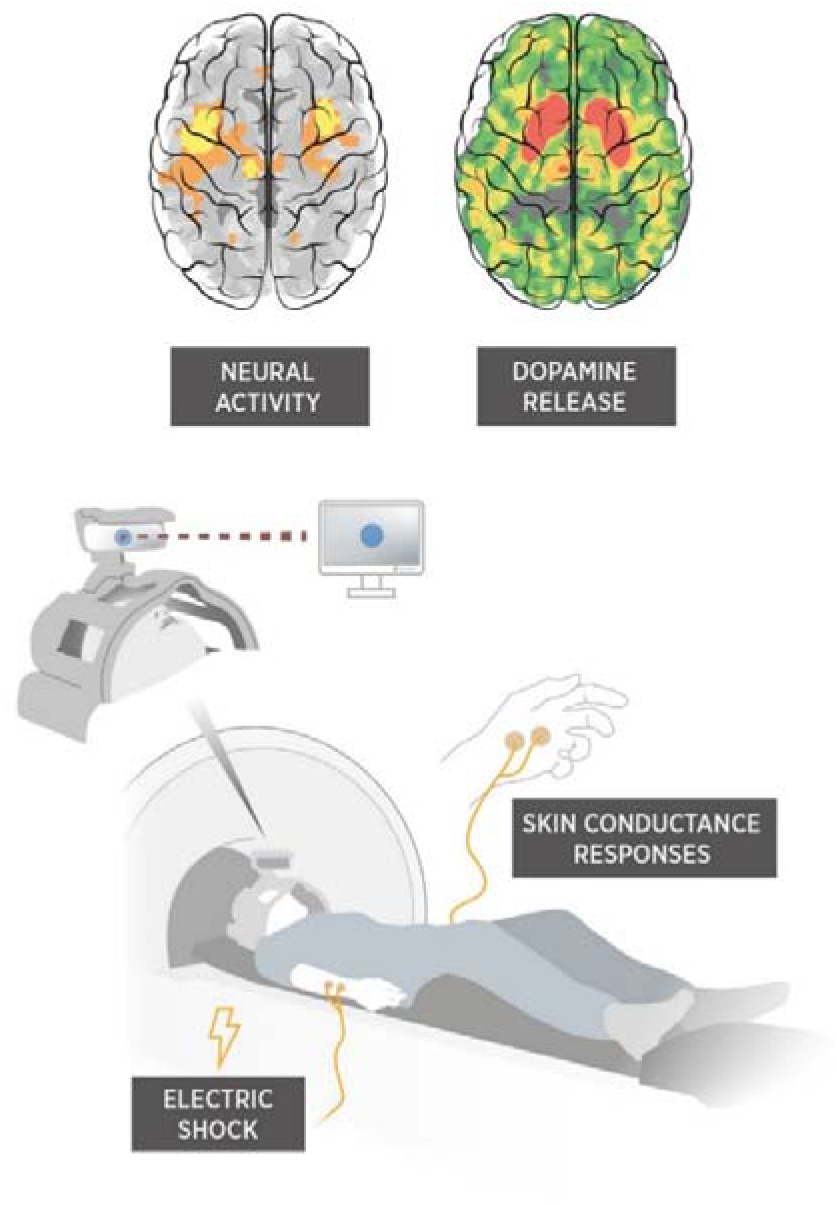
Simultaneous measures of dopamine release and neural activity in the amygdala during fear conditioning probed with standard skin-conductance response. Positron emission tomography using single scan bolus plus constant infusion of [^11^C]raclopride was combined with event-related functional magnetic resonance imaging during fear conditioning, where one cue displayed on the screen was reinforced by a mild electric shock (CS+), while another control cue (CS-) never was paired with shock. Brain images for illustration only and do not depict the actual results of the study.

## MATERIALS AND METHODS

### Experimental Design

Eighteen individuals (mean ± SD age 25.2 ± 4.8 years; 10 women, 8 men; 16 right-handed, 2 left-handed) recruited through local advertisements were included in the study, which was approved by the regional ethics review board and radiation safety committee in Uppsala, Sweden. Participants arrived at the scanning site about 2 hours before scanning. They were informed about the study and signed informed consent. Approximately 90 minutes prior to radiotracer injection and PET scanning, participants determined the strength of the unconditioned electric shock through a staircase procedure with the instruction that the shock should be unpleasant, but endurable.

Participants were positioned supine in the combined Signa 3T PET/MR scanner (GE Healthcare) with their heads lightly fixated inside the head coil. A bolus (20 s) of the selective dopamine D2/3 receptor antagonist [^11^C]raclopride was injected through a venous catheter and followed by constant infusion during the 90 minutes of PET data acquisition. Following 50 minutes of resting PET data collection, participants underwent a differential fear conditioning paradigm during collection of blood-oxygenation-level dependent (BOLD) fMRI and skin conductance. The PET scanning continued 20 minutes after the fear conditioning paradigm.

### Fear Conditioning Paradigm

The fear conditioning paradigm lasted 20 minutes and consisted of 20 presentations each of two geometrical shapes (a brown arrow and a blue circle) used as conditioned stimuli (CS). One of the shapes (CS+) was paired with an electric shock on 16 occasions (80% reinforcement rate), and the other (CS-) was unpaired<u>(*23*)</u>. The CSs were counterbalanced across subjects. Each CS was presented for 6 s with a mean 24.3 s fixation cross inter-trial interval varying between 21.8 and 27 s. CS+ co-terminated with a 250 ms electric shock on reinforced trials.

Visual stimuli were projected onto a 32” computer screen positioned at the head of the scanner using E-prime 2 (Psychology Software Tools, Pittsburgh, PA, USA). Participants viewed the computer screen through a mirror on the head-coil. The presentation software was synced with fMRI data acquisition using a SyncBox (NordicNeuroLab, Bergen, Norway).

Electric shocks were used as unconditioned stimuli (US) and delivered to the subjects’ dorsal right lower arm via disposable radiotransluscent electrodes (EL509, BIOPAC Systems, Goleta, CA, USA) by the STM100C module connected to the STM200 constant voltage stimulator and controlled by the BIOPAC MP150 (BIOPAC Systems, Goleta, CA, USA).

### Skin Conductance Responses

Skin conductance was recorded using the BIOPAC MP150 (BIOPAC Systems, Goleta, CA, USA). Disposable radiotransluscent Ag/AgCl electrodes (EL509 Biopac electrodes) were filled with isotonic electrode gel (GEL101 Biopac gel) and applied to the hypothenar eminence of the left hand. A 0.05 Hz high-pass filter was applied to the signal. SCRs were calculated as the maximum phasic driver amplitude 1-5 s after stimulus presentation using the Ledalab software(*24*) implemented in MATLAB (Mathworks Inc., Natick, MA) with responses <0.01 scored as 0 (non-response). SCRs were square root transformed and range-corrected by dividing each participant’s SCRs with his/her maximum SCR, resulting in SCRs ranging from 0 to 1. This minimizes the influence of individual differences and isolates the experimental effects.

To evaluate fear conditioning acquisition, mean values of SCRs to CS+ and CS-were used to calculate delta scores (CS+ minus CS-) for each individual. Delta scores are independent of individual differences in general reactivity and habituation (response decline over successive trial presentations) as that affects CS+ and CS-to an equal extent and thus control for nonspecific activity and represents an unbiased measure of associative learning with respect to general arousal. All methods are standard procedures and analytic strategies in fear conditioning(*25, 26*).

### PET/fMRI Acquisition

Subjects underwent a 90 min PET scan on a 3T Signa PET-MR scanner (GE Healthcare, Waukesha), initiated simultaneously with the start of a [^11^C]raclopride bolus-infusion protocol (total amount of radioactivity mean±SD 379±75 MBq; kbol 107 min). Images were reconstructed into 18 5-min frames using ordered subset expectation maximization (4 iterations, 28 subsets), including resolution recovery and a 5 mm Gaussian post-filter. Attenuation correction was done using the manufacturer’s atlas-based method, and all other corrections necessary for quantitative PET images were applied. Atlas-based attenuation correction has been shown to be less accurate than CT-based or zero echo time (ZTE) MRI-based attenuation correction, but because we were only addressing changes in receptor binding within the same patient and scan, this does not affect our results. Head movement in the scanner was restricted using foam cushions.

Anatomical 3D T1-weighted images were acquired with an 8 channel head coil and the following parameters (repetition time (TR)=8.6 ms, echo time (TE)=3.3 ms, inversion time=450 ms, flip angle=12, matrix=256×256, voxel size=1.2×1.2×1.2 mm), starting approximately 15 minutes after bolus injection. BOLD fMRI was collected using a single shot echo-planar imaging (EPI) sequence with parameters (TR=3000 ms, TE=30 ms, flip angle 90°, matrix=64×64, voxel size=3.0×3.0 mm, slice gap=0.4 mm, slices=45).

### PET Analysis

PET images were corrected for inter-frame motion using frame-by-frame alignment with Voiager software (GE Healthcare, Uppsala, Sweden). For each participant, the T1-weighted MRI image was co-registered to a summed PET image and segmented into gray matter, white matter and CSF using SPM8 (fil.ion.ucl.ac.uk/spm). A probabilistic volume of interest (VOI) template containing 45 VOIs was applied to the co-registered T1-weighted MRI image and transferred to the dynamic PET data using PVElab (*27*), resulting in gray matter time-activity curves (TAC) of the regions of interest.

PET data were analyzed both in VOIs and voxel-by-voxel using a nested two-step approach: first, the initial 50 min of the TACs were analyzed using the simplified reference tissue model (SRTM) (*28*) for VOIs and for voxel-wise analyses a basis function implementation of SRTM (*29*) with cerebellar gray matter as reference tissue, giving baseline R_1_ (tracer delivery relative to cerebellum), k’_2_ (reference tissue efflux rate constant) and BP_ND_ (non-displaceable binding potential). Then, the same models were applied to the 70-90 min interval, fixing R_1_ and k’_2_ at baseline values and only fitting for BP _ND_, the binding potential after conditioning. The percent difference in binding potential was calculated as [100 × (1-BP _ND_/BP_ND_)] and used as a measure of endogenous dopamine release. The analyses resulted in VOI values and parametric images of BP_ND_, BP’_ND_ and dopamine release. The parametric images were normalized to Montreal Neurological Institute (MNI) standard space in SPM12 by first coregistering each individual’s parametric images to their T1-weighted MRI image, then segmenting and normalizing the T1-weighted image to MNI space, and finally applying the transformation parameters to the PET images. Images were resliced to 4 mm isotropic voxels and smoothed with an 8 mm full width half maximum (FWHM) Gaussian kernel.

An illustration of the fit of the two SRTM models to mean TACs is presented in Fig. 2. If there would be no dopamine release (i.e. no change in [^11^C]raclopride BP_ND_), the time-activity data would follow the fit line of the 0-50 minute data. As can be seen in Fig. 2, the time-activity data in the amygdala and striatum no longer follow the expected curve from the start of fear conditioning at 50 min post injection, indicating dopamine release resulting in a reduced BP_ND_. This deviation cannot be seen in the frontal cortex, thus not indicating dopamine release in this region.

**Fig. 2.**
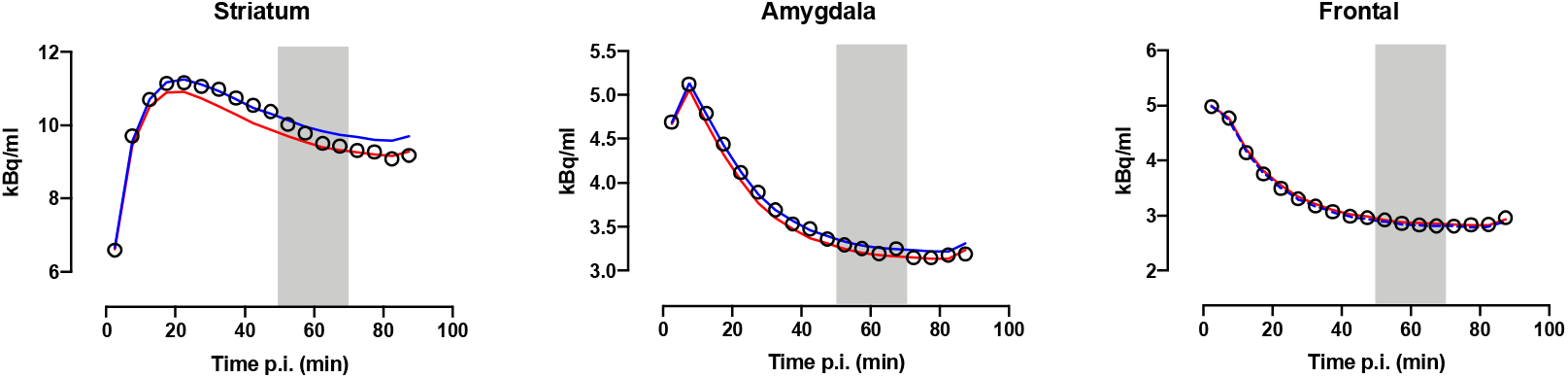
Simplified reference tissue model fits. Mean time-activity curves in the striatum, amygdala, and frontal cortex. Time post injection (p.i.) minutes. Blue lines are fit lines of the simplified reference tissue model (SRTM) fits to the 0-50 min baseline portion of the data, whereas red lines are fits to the 70-90 min post conditioning portion of the data re-using all parameters from the baseline fit except BP_ND_. The gray square denotes the timing of the fear conditioning at 50-70 min p.i. If there would be no dopamine release, i.e. no change in [^11^C]raclopride binding potential (BP_ND_), the time-activity data would follow the fit line of the 0-50 minute data. As can be seen, the PET time-activity data in the amygdala and striatum no longer follow the expected (blue) curve from the start of fear conditioning at 50 min post injection, indicating dopamine release resulting in a reduced BP_ND_. The fits to the mean time-activity curves correspond to a reduction in BP_ND_ after challenge of 5.8%, 12.5%, and 0% in the striatum, amygdala, and frontal cortex, respectively.

Ideally, the use of a bolus-infusion protocol results in a steady state, which would allow for use of simple radioactivity concentration ratios to measure occupancy instead of tracer kinetic modelling. However, the nested two-step SRTM method can account for deviations from steady state, which are nearly always present, whilst ratio methods cannot. Another, previously published analysis method (lp-ntPET) (*30*) would have given us the time course of the dopamine release in addition to the magnitude. This method, requiring fitting of 6 parameters instead of the 4 parameters in the present work, is less robust because of the larger number of parameters. Since neurotransmitter release due to fear conditioning is likely of lower levels than that due to pharmacological challenges, robustness of the model to measure small changes in BP is necessary.

Robustness of measuring post-challenge BP_ND_ was assessed by performing a simulation study. For dopamine release levels resulting in a reduction in BP_ND_ of 0, 5, 10, 15 and 20%, 100 time-activity curves were numerically simulated using published values of rate constants, typical noise levels, and baseline BP_ND_ values of 2.6 (representing striatum), 0.3 (representing amygdala), and 0.07 (representing frontal cortex). Changes in BP_ND_ during the scan were simulated by a linear reduction in k_3_ between 50 and 70 min post injection, and resulting time-activity curves were analyzed using the nested version of SRTM. In addition, 100 TACs with a range of BP_ND_ changes between 0 and 30% were simulated for each of the baseline BP_ND_ values. A minor bias in post fear conditioning BP_ND_ of 0-2% and coefficient of variation (COV) of around 10% was found for baseline BP_ND_ values of 0.3 (amygdala). For high baseline BP_ND_ (striatum), there was a positive bias in post-challenge BP_ND_ that was proportional to dopamine release levels, and a COV of around 2.5%. For lower baseline BP_ND_ (frontal cortex), a varying bias of ±5% was seen with COV exceeding 40%. Hence, for striatum, the nested SRTM method seems to result in a proportional underestimation of dopamine release, whereas this effect is much smaller for amygdala (Fig. 3 and 4). Postchallenge BP_ND_ for frontal cortex cannot be estimated reliably because of poor precision. Use of SUVR or lp-ntPET resulted in considerably larger bias and variability (data not shown).

**Fig 3.**
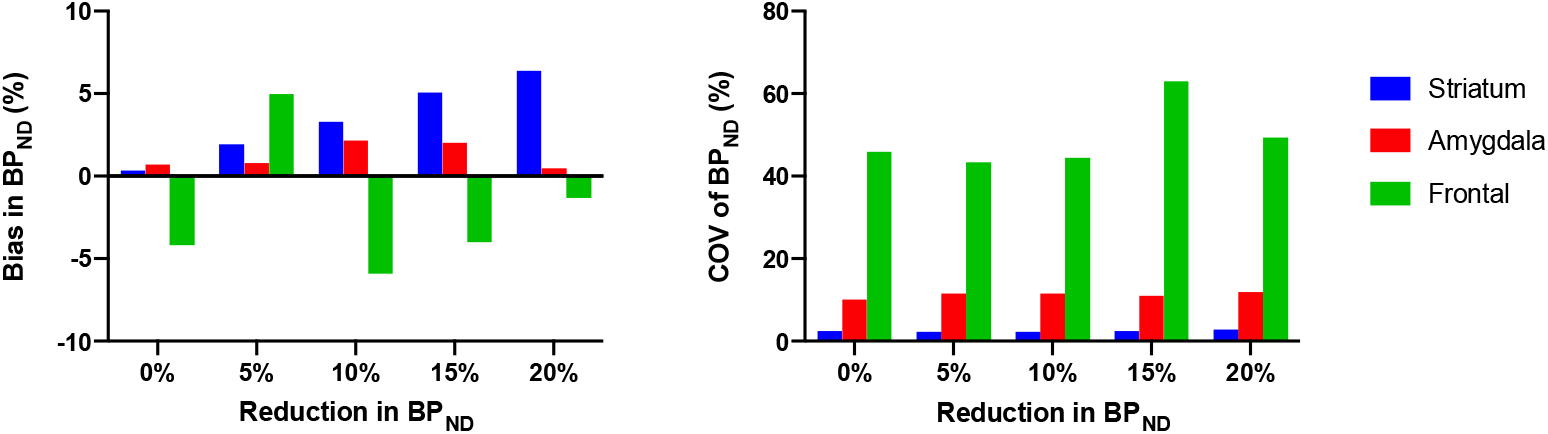
Bias and coefficient of variation of measures of post fear conditioning binding potential. Bias (left) and coefficient of variation (COV; right) of binding potential (BP_ND_) values post fear conditioning based on 100 simulated time-activity curves resulting in 0-20% decrease in receptor availability, for baseline BP_ND_ of 2.6 (representing striatum), 0.3 (representing amygdala), and 0.07 (representing frontal cortex). A minor bias in post-challenge BP_ND_ of 0-2% and COV of around 10% was found for baseline BP_ND_ values of 0.3. For high baseline BP_ND_, there was a positive bias in postchallenge BP_ND_ that was proportional to dopamine release levels, and a COV of around 2.5%. For lower baseline BP_ND_, a varying bias of ±5% was seen with COV exceeding 40%.

**Fig. 4.**
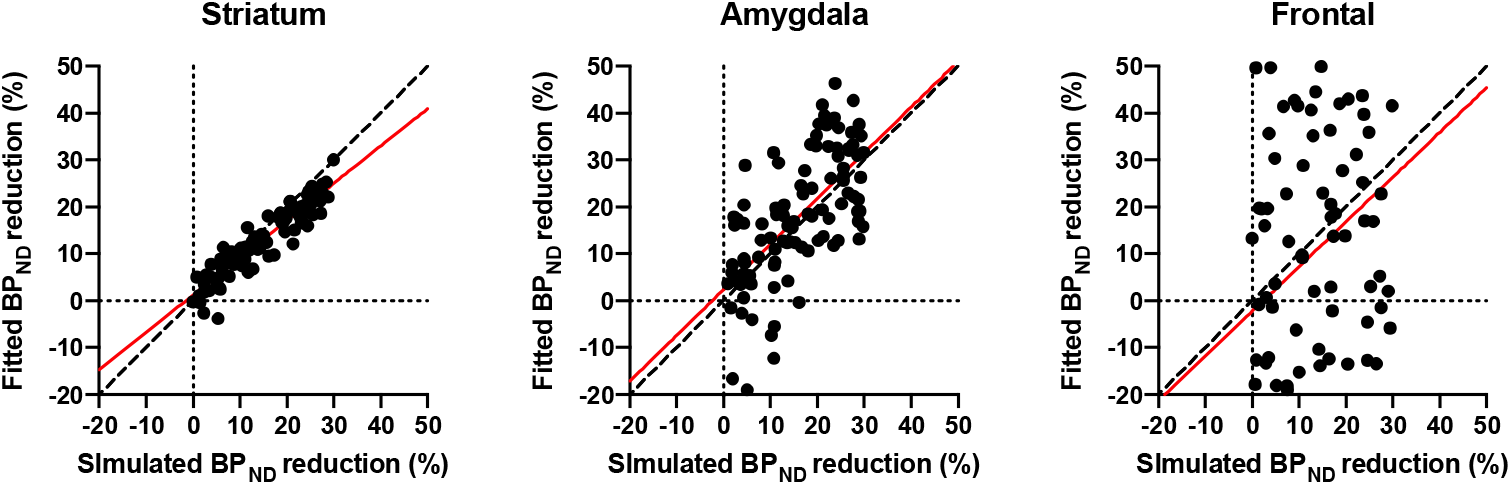
Fitted binding potential reduction versus simulated binding potential reduction. Binding potentials (BP_ND_) were fitted and change in BP_ND_ calculated for 100 simulated time-activity curves with 0-30% reduction in BP_ND_. Plots show fitted BP_ND_ reduction plotted against the simulated BP_ND_ reduction for the striatum, amygdala, and frontal cortex. Dashed lines represent the identity line and red solid lines the linear fit line. For baseline BP_ND_ of 2.6 (striatum), there was a negative bias in fitted BP_ND_ reduction that was proportional to dopamine release levels, in line with the positive bias in fitting post fear conditioning BP_ND_ (Fig. 3). For baseline BP_ND_ values of 0.3 (amygdala), there was a minor bias, whereas for BP_ND_ of 0.07 (frontal cortex), the scatter plot clearly illustrates the lack of reliable fit of BP_ND_ reductions at these levels.

Thus, simulations show a high accuracy and precision of the two-step nested method applied in the present work provided that the dopamine release time course, modelled as a gamma variate function, peaks before 70 min and has a relatively slow reduction after that. These conditions were confirmed both by microdialysis experiments in rodents(*31*) and by the observation that we see a continually reduced signal after 70 min in our data, with no reversal to initial activity concentration values within the time course of the scans, suggesting a persisting change in dopamine levels. Furthermore, simulations showed that the occupancy values estimated by this model were insensitive to conditioning-induced changes in R_1_.

### fMRI Analysis

FMRI data was preprocessed and modeled using SPM12 (fil.ion.ucl.ac.uk/spm/software/spm12) implemented in MATLAB R2018a (Mathworks Inc., Natick, MA). Each individual’s fMRI data was first slice-timing corrected, realigned and coregistered to the anatomical T1-weighted image, and then normalized to isotropic 4 mm voxels in the MNI standard space by applying the transformation parameters from the segmentation of the T1-weighted image. Finally, the images were smoothed with an 8 mm FWHM Gaussian filter.

The first level model for each participant was fitted with onsets and durations of CS+, CS-, and shock (modeled as a stick function) convolved with the canonical haemodynamic response function from SPM12, together with the six realignment parameters from the realignment step, as regressors. Contrast images were created for CS+, CS- and US vs. baseline, and for the learning-related CS+ minus CS-. Mean BOLD responses within the regions of interest amygdala and striatum were extracted using the Wake Forest University School of Medicine (WFU) PickAtlas automatic anatomical labeling (AAL) definitions of these regions (*32*).

To address potential influence of motion on the results, in addition to including the realignment parameters in the first level model, we set a cut-off to keep subjects with head motion not exceeding one acquisition voxel (i.e., 3 mm) in any direction (*33*). We also calculated the total net movement during the fear conditioning task (mean ± SD: 1.3 ± 0.7 mm) and examined the relation between this measure and change in [^11^C]raclopride BP_ND_, but could not detect any association in the amygdala (*r*=-0.01, *p*=0.960, 95% CI: −0.48 to 0.46) or in the striatum VOIs (*r*=0.24, *p*=0.343, 95% CI: −0.26 to 0.63).

### Statistical Analyses

We applied three a priori regions of interest in our analyses, the amygdala, striatum, and frontal cortex (superior frontal gyrus). The amygdala was our primary target based on our hypotheses, the striatum (combined caudate nucleus, putamen and nucleus accumbens) was included based on the involvement in fear conditioning and rich dopamine receptor D2/3 expression in this region (*22*), and the frontal cortex (superior frontal gyrus), low in D2/3 receptor expression was our control region where we expected not to be able to measure dopamine release robustly.

Our main analyses involved the [^11^C]raclopride BP_ND_ VOI values and mean BOLD responses within the amygdala and striatum. We also applied voxel-wise paired t-tests (baseline vs postconditioning) in SPM12 to locate overlapping voxels with both lower [^11^C]raclopride BP_ND_ (i.e., dopamine release) and learning-related neural activity (CS+ minus CS-BOLD responses) from fear conditioning. The statistical threshold was set to P<0.05 family-wise (FWE) corrected for multiple comparisons using random field theory.

To assess the relations between dopamine release and central and peripheral learning, and between central and peripheral learning, we entered dopamine release and neural activity from the amygdala and striatum ROIs into Pearson’s product-moment correlations together with the learning-related delta SCR (CS+ minus CS-) in R version 4.0.0 (*34*). We used directed tests (P<.05) due to the a priori hypotheses of positive correlations between these measures.

## RESULTS

### Dopamine release during fear conditioning

First, using single scan bolus/infusion [^11^C]raclopride PET we found decreased binding potential (BP_ND_) from baseline to post conditioning in the amygdala (Fig. 5A), confirming the primary hypothesis of dopamine release in this region during fear conditioning. Additionally, exploratory analyses revealed decreased BP_ND_ following conditioning in the striatum (Fig. 5B), indicating dopamine release also in this brain area. In the frontal cortex, included here as a control region where we expected no decrease in BP_ND_, we could not detect any change in BP_ND_ between baseline and post fear conditioning (mean change: −34.6%, 95% CI: −159.8% to 90.5%) (*t*(17)=0.54, *p*=0.595). Complementing these analyses of mean dopamine release in each region, we performed within region voxel-wise analyses, revealing reduced BP_ND_ in bilateral amygdala clusters (Fig. 6) and in the striatum (Fig. 7).

**Fig. 5.**
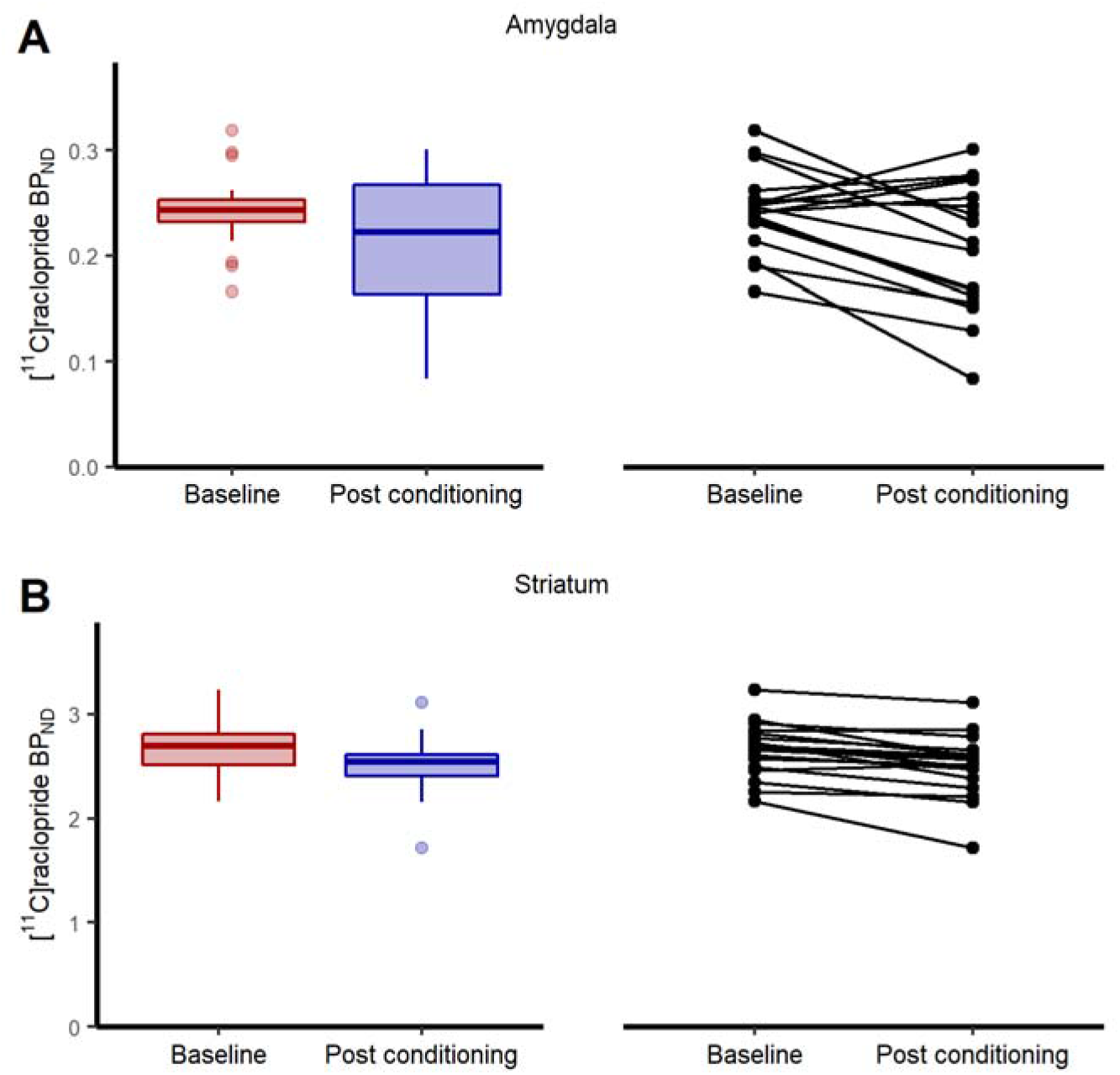
Binding potential of [^11^C]raclopride at baseline and after fear conditioning. Boxplots and individual participant’s trajectory lines showing [^11^C]raclopride binding potential (BP_ND_) at baseline and after fear conditioning in anatomically defined regions of interest. **(A)** In accordance with our hypothesis, [^11^C]raclopride BP_ND_ in the amygdala decreased by 13.4% (95% CI: 3.8% to 22.9%) (*t*(17)=2.74, *p*=0.007) from baseline to after fear conditioning, indicating dopamine release. **(B)** Likewise, in the striatum, there was a 5.9% (95% CI: 3.4% to 8.4%) decrease in BP_ND_ (*t*(17)=4.69, *p*=0.0002). For the boxplots, the line indicates the median, the box the interquartile range (IQR), the whiskers the minimum of 1.5×IQR and minimum/maximum values, and circles values more extreme than 1.5×IQR. Data for individual participants is shown in the trajectory lineplots.

**Fig. 6.**
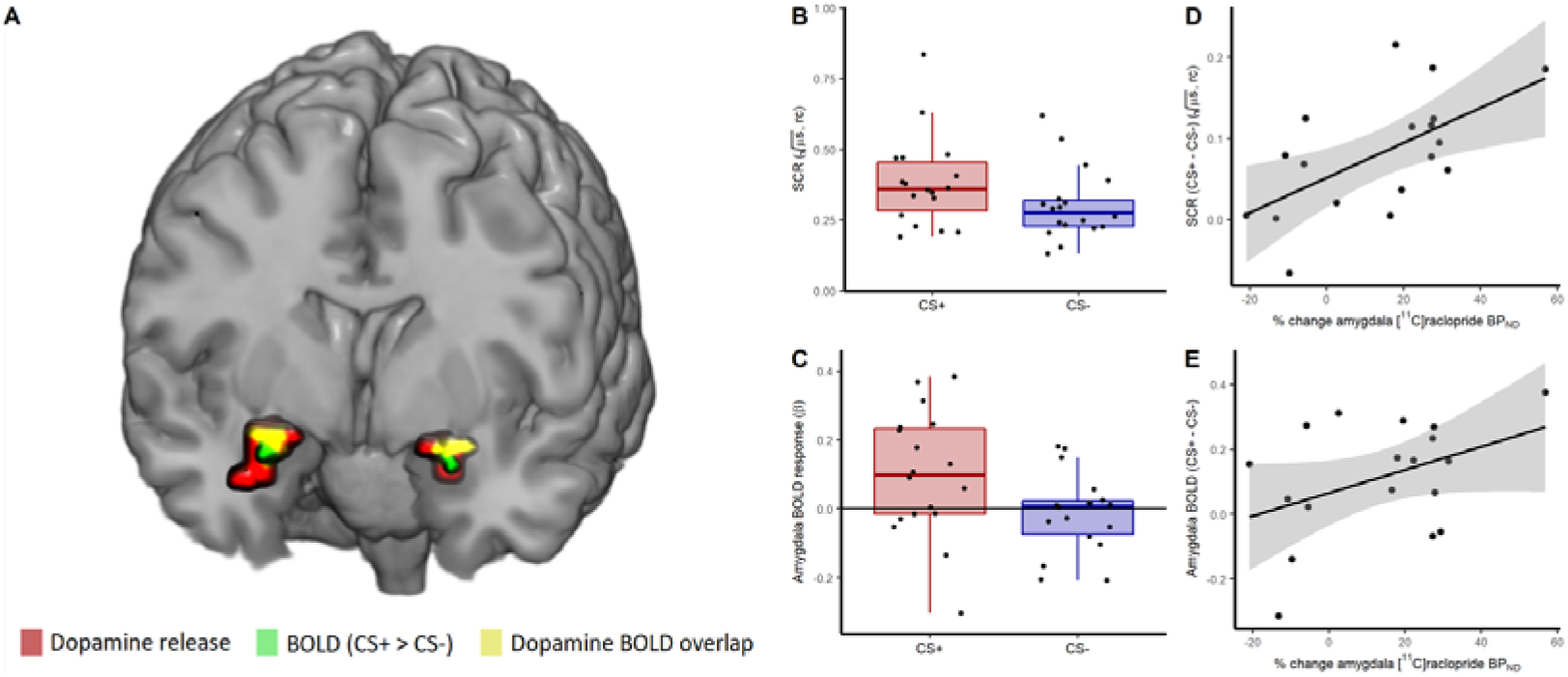
Dopamine release in the amygdala during fear conditioning was co-localized with the neural memory trace and predicted both learning strength and memory trace activity. **(A)** [^11^C]raclopride binding potential (BP_ND_) was decreased during fear conditioning in a 320 mm^3^ volume in the left (Z=3.22 P_FWE_=.017 family-wise error [FWE] corrected) and a 960 mm^3^ volume in the right amygdala (Z=3.35; P_FWE_=.012) with peak voxels at Montreal Neurological Institute coordinates −22, 0, −14 and 26, 0, −14 respectively, indicating dopamine release in these areas, and co-localized with bloodoxygenation-level dependent (BOLD) response to conditioned stimuli (CS+ - CS-) in the left (Z=3.44 P_FWE_=0.009; MNI −22, 0, −14) and right amygdala (Z=3.04; P_FWE_=0.028; MNI: 22, 0, −14), indicating memory formation. Red clusters show dopamine release, green denotes clusters with greater neural activity to CS+ than CS-shown here thresholded at *p*<0.05 for illustrative purposes, and yellow signifies overlap between dopamine release and learning-related neural activity within the amygdala. The fear conditioning procedure resulted in **(B)** discrimination between fear (CS+) and control (CS-) cues on skin conductance responses (SCR) (*t*(17)=4.64, *p*=0.0001) and **(C)** in an amygdala-located memory trace (CS+ > CS-) (*t*(17)=2.70, *p*=0.008). The amount of dopamine release in the amygdala predicted **(D)** learning strength (*r*(16)=0.60, *p*=0.004, 95% CI: 0.27 to 1) and **(E)** amygdala memory trace activity (*r*(16)=0.41, *p*=0.044, 95% CI: 0.02 to 1). For the boxplots, the horizontal line indicates the median, the box the interquartile range (IQR), the whiskers the minimum of 1.5×IQR and minimum/maximum values, and the filled circles data for individual participants. For the scatterplots, shaded areas reflect standard error of means. rc: range corrected to each individual’s maximum SCR. All statistical tests are one-sided tests of directed hypotheses.

**Fig. 7.**
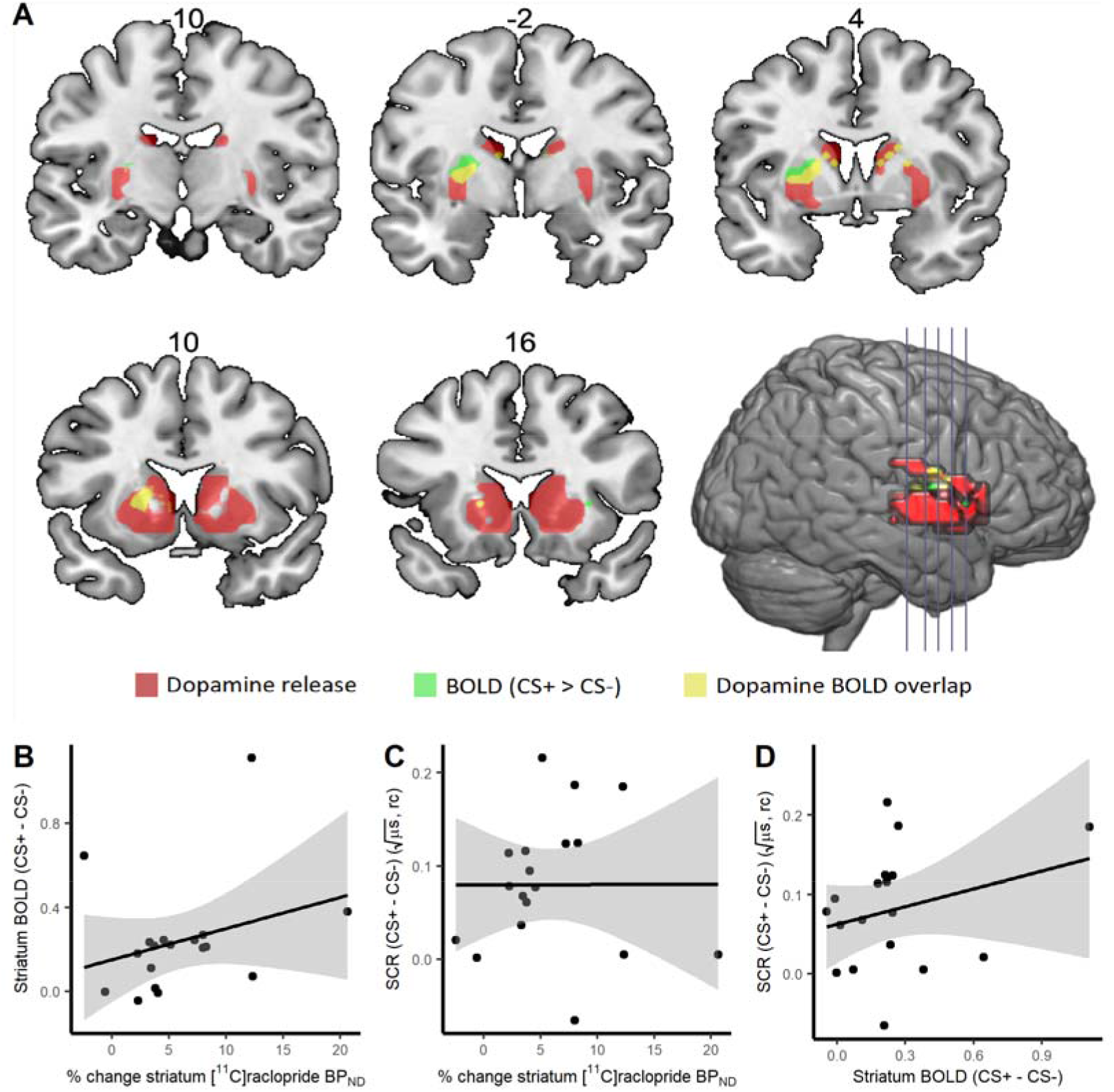
Striatal dopamine release, neural activity and skin conductance response during fear conditioning. **(A)** [^11^C]raclopride binding potential (BP_ND_) was decreased during fear conditioning in a 7488 mm^3^ volume in the left (Z=6.24 P_FWE_<.001 family-wise error [FWE] corrected) and a 8320 mm^3^ volume in the right striatum (Z=6.54; P_FWE_<.001) with peak voxels at Montreal Neurological Institute coordinates −18, 8, −2 and 22, 8, −2 respectively, indicating dopamine release in these areas, and colocalized with blood-oxygenation-level dependent (BOLD) response to conditioned stimuli (CS+ - CS-) in the left (Z=4.87 P_FWE_=0.0002; MNI −22, 0, 6) and right dorsal striatum (Z=4.07; P_FWE_=0.008; MNI: 22, 0, 10), indicating memory formation. Red clusters show dopamine release, green denotes clusters with greater neural activity to CS+ than CS-, and yellow signifies overlap between dopamine release and learning-related neural activity within the amygdala. We could not detect any relation between percent change in striatal [^11^C]raclopride BP_ND_, i.e. dopamine release, and **(B)** blood-oxygenation-level dependent (BOLD) response to conditioned stimuli (CS+ - CS-) (*r*(16)=0.29, *p*=0.237, 95% CI: −0.20 to 0.67), or **(C)** the peripheral measure of fear learning, skin conductance responses (SCR) (*r*(16)=0.003, *p*=0.991, 95% CI: −0.46 to 0.47). **(D)** Nor could we detect a relation between striatal BOLD response and SCR (*r*(16)=0.27, *p*=0.271, 95% CI: −0.22 to 0.66]. rc: range corrected to each individual’s maximum SCR. For the scatter plots, shaded areas reflect standard error of means.

### Fear conditioning procedure induced learning

Second, fear conditioning resulted in skin conductance response discrimination between fear and safety-predicting cues (CS+ > CS-), i.e. a peripheral expression of learning, as evidenced by the repeated measures ANOVA revealing main effects of CS (*F*(1, 17)=21.53, *p*=0.0002) and Trial (*F*(19, 323)=23.96, *p*<0.00001), and a CS × Trial interaction (*F*(19, 323)=2.249, *p*=0.0023) (Fig. 6B). Blood-oxygenation-level dependent (BOLD) fMRI revealed an amygdala-localized memory trace (i.e. BOLD response CS+ > CS-) (Fig. 6A, 6C) that was linearly coupled to the skin conductance responses (*r*(16)=0.44, *p*=0.033, 95% CI: 0.05 to 1.00), consistent with a vast literature underscoring amygdala as a key brain structure for aversive memory formation in humans and other animals (*1*).

### Dopaminergic facilitation of fear learning

Next, we tested if dopamine release and learning strength are functionally coupled in the amygdala by correlating percent change in [^11^C]raclopride BP_ND_ with SCR difference scores (CS+ - CS-) and found a positive linear relationship (Fig. 6D). In contrast, the unconditioned response to electric shocks was not related to dopamine release in this region (*r*(16)=0.31, 95% CI: [-0.18 to 0.68], *p*=0.209]). Percent change in amygdala [^11^C]raclopride BP_ND_ also predicted learning-induced neural activity in the amygdala (Fig. 6E). These findings confirm dose-response relations between amygdala dopamine release and learning strength, both in the peripheral and central nervous systems. Areas in the amygdala with endogenous dopamine release overlapped with the areas reflecting neural memory trace activity (Fig. 6A), demonstrating that dopamine release and neural activity were both functionally and anatomically coupled. Also, in the dorsal striatum, there was spatial overlap between dopamine release and neural activity (CS+ > CS-) (Fig. 7A). However, we could not detect any relations between striatal dopamine release and striatal BOLD response (Fig. 7B) or SCR during conditioning (Fig. 7C), nor between striatal BOLD response and SCR (Fig. 7D), indicating specificity of dopaminergic facilitation of memory formation in the amygdala.

## DISCUSSION

We show that human fear conditioning is associated with endogenous dopamine release in the amygdala and that learning strength changes in concert with dopamine release. This mirrors rodent studies demonstrating that fear conditioning is dependent on dopamine signaling in the amygdala (*9-11, 13, 14, 35-37*). Using strict experimental controls, we confirm that fear conditioning induces peripheral and central nervous system learning. Statistically, we could further demonstrate that dopamine facilitates fear learning since learning strength was linked to dopamine release, while the strength of the unconditioned reaction was not. This is consistent with an interpretation that dopamine drives or is driven by learning-related processes and suggesting specificity for learning-induced processes over stress reactivity. The association between dopamine and conditioned, but not unconditioned, responses is consistent with a study in rodents reporting that fear conditioning induced a higher rise in dopamine concentration than did shock presentations only (*38*). Two recent rodent studies have also showed that dopamine is released during foot shock and necessary for fear conditioning (*37*), and that the same dopaminergic neurons projecting from the ventral tegmental area to the basal amygdala are activated by aversive and appetitive stimuli as well as cues that predict the aversive and appetitive stimuli (*39*). These findings would rather point to the role of dopamine signaling salience.

One possible mechanism explaining our results is that dopamine facilitates long-term potentiation (LTP). LTP in the amygdala can cause fear learning (*40*), and because dopamine release facilitates LTP (*41*), our data are consistent with the notion that fear learning in humans is facilitated by dopamine-induced LTP and also with a recent rodent study demonstrating that D2 receptor stimulation facilitates fear learning (*42*). We suggest that dopamine serves as a neurochemical guide to strengthen aversive memory formation and behavioral output.

Furthermore, there was evidence of an overlap between dopamine release in the striatum and learning-related activity. The voxel-wise analyses indicated overlap between PET and fMRI measures in the dorsal striatum, but the functional implications of this overlap is unclear as we could not detect any correlation between the two, or between these measures and the peripheral learning index. We can only speculate that the striatal dopamine release may be related to avoidance action programs.

Limitations of the current study deserve mentioning. First, the use of [^11^C]raclopride to measure dopamine release outside the striatum has been questioned. This is based on the lack of change in BP in pharmacological occupancy studies. However, these studies have rarely included the amygdala or applied single scan bolus/infusion paradigms which arguably adds to the sensitivity of the measure. Moreover, test-retest reliability of amygdala [^11^C]raclopride BP has been reported to be high (*21*). Also, here we found similar changes in [^11^C]raclopride BP_ND_ in the amygdala and the striatum, although amygdala release had higher variability, whereas our control region, the frontal cortex did not indicate dopamine release. In addition, simulations (Fig. 3, 4) indicated that reduction in [^11^C]raclopride binding potential can be robustly measured in the amygdala using the nested two-step simplified reference tissue model (SRTM) approach used in this study.

Using simultaneous measures of dopamine release and neural activity in a combined PET/MRI scanner we show that dopamine release during fear conditioning is linked to synaptic plasticity and aversive memory formation. This suggests that blocking dopaminergic signaling would reduce fear memory acquisition and that augmenting dopamine transmission during memory extinction would strengthen safety memory formation. This has clinical implications as manipulating fear and safety memories through extinction-based exposure forms the basis for cognitive behavioral therapy in anxiety and post-traumatic stress disorders (*43*). Consistently, dopamine infusions in the rodent basolateral amygdala facilitate memory consolidation (*44*) and in humans, systemic L-DOPA administration strengthens safety memories during experimental fear extinction (*45*). Dopamine facilitates memory formation, not only in the neural fear circuitry, but also in non-associative learning (*46*), instrumental conditioning (*5*), and in cognitive episodic memory (*47*). Thus, we argue that dopamine represents an evolutionary conserved neurochemical mechanism supporting learning across multiple memory systems.

## Acknowledgments

We thank the staff at the Uppsala PET/MR facility and Uppsala PET Center for invaluable help with data collection.

## Author contributions

A.F.: Methodology, Formal analysis, Investigation, Data curation, Writing – Review & Editing, Visualization, Project administration, Funding acquisition; J.B.: Methodology, Investigation, Writing – Review & Editing; M.L.: Methodology, Software, Data curation, Supervision, Writing – Review & Editing; A.E.: Investigation, Writing – Review & Editing; M.F.: Conceptualization, Methodology, Writing – Original Draft, Supervision, Funding acquisition; F.Å.: Conceptualization, Methodology, Writing – Review & Editing, Funding acquisition.

## Data and materials availability

The data that support the findings of this study are available from the corresponding author upon reasonable request.

